# Construction of a novel phagemid to produce custom DNA origami scaffolds

**DOI:** 10.1101/309682

**Authors:** Parsa M. Nafisi, Tural Aksel, Shawn M. Douglas

**Affiliations:** Department of Cellular and Molecular Pharmacology, University of California, San Francisco, San Francisco, CA 94158, USA

## Abstract

DNA origami, a method for constructing nanoscale objects, relies on a long single strand of DNA to act as the “scaffold” to template assembly of numerous short DNA oligonucleotide “staples”. The ability to generate custom scaffold sequences can greatly benefit DNA origami design processes. Custom scaffold sequences can provide better control of the overall size of the final object and better control of low-level structural details, such as locations of specific base pairs within an object. Filamentous bacteriophages and related phagemids can work well as sources of custom scaffold DNA. However, scaffolds derived from phages require inclusion of multi-kilobase DNA sequences in order to grow in host bacteria, and thus cannot be altered or removed. These fixed-sequence regions constrain the design possibilities of DNA origami. Here we report the construction of a novel phagemid, pScaf, to produce scaffolds that have a custom sequence with a much smaller fixed region of only 381 bases. We used pScaf to generate new scaffolds ranging in size from 1,512 to 10,080 bases and demonstrated their use in various DNA origami shapes and assemblies. We anticipate our pScaf phagemid will enhance development of the DNA origami method and its future applications.

## Introduction

DNA origami builds tiny shapes using long single-stranded DNA (ssDNA) scaffolds and short ssDNA staples (1). A key strength of this method is that one scaffold sequence can be reused with different staples to create many different shapes. However, although shapes generated from a single scaffold sequence can vary in nanometer-scale geometry, it is difficult to engineer sub-nanometer-scale details of shapes using only a single scaffold, or to create structures that vary in size from a single scaffold precursor (2). Thus, it is important to generate new scaffolds to expand the space of possible DNA origami designs. For example, new scaffolds could help create larger structures, or shapes with novel multimeric assemblies, or could help elucidate the principles of DNA origami self-assembly. Here, our goal was to create new scaffolds of almost arbitrary custom sequence, up to 10 kilobases (kb) long, which are suitable for production in milligram (mg) quantities.

Many methods for producing scaffolds of custom sequence have been reported previously (3–17). Typically, ssDNA scaffolds are derived from double-stranded DNA (dsDNA) sources via combinations of selective amplification, isolation, or degradation of one of the two strands. While reported methods offer excellent sequence customizability, they can be difficult to scale up in sequence length and production yield due to limitations in various *in vitro* enzymatic processing steps. We set out to develop a general approach for easily generating new scaffolds that would be scalable in both length and production yield, thus overcoming a significant hurdle in expanding their usability.

We selected filamentous bacteriophages as a platform that offers mg-scale yields in shake flasks, and whose yield can be boosted with bioreactors if needed (20). Filamentous phages, such as M13, are bacteria-specific viruses that package and export their single-stranded genomes into rod-like particles that have a protein coat. The phages can be recovered from the culture media and the ssDNA purified by standard molecular biology techniques (21). Custom sequences up to 2.5 kb can be inserted reliably into the M13 genome (22), though schemes to create much longer scaffolds have been reported (13). However, sequence customizability in the M13 phage is limited because most of its genome (>6 kb) consists of protein-coding and regulatory sequences that cannot be easily modified without disrupting phage growth.

Phagemids can be used to create scaffolds that have improved sequence customizability compared to M13 (7, 15, 23). These plasmids typically contain a host origin of replication (ori) sequence, a phage ori from M13 or relative such as f1, and an antibiotic resistance gene. Phagemid ssDNA can be exported in phage-like particles if the host cell is co-infected with a “helper phage” or transformed with a “helper plasmid” to express the necessary viral proteins (24). Phagemids can accommodate custom inserts several kb in size, but include a 2–3 kb fixed region that limits their usefulness in producing custom origami scaffolds.

To increase phagemid sequence customizability, we sought to create a scaffold that could be produced using established preparation methods but would be able to package and export custom ssDNA sequences that have a relatively small fixed region. We were inspired by two papers that reported making use of modified f1-ori sequences to manipulate ssDNA synthesis (Fig. 1 D and E). In 1982, Dotto *et al*. used phagemids with modified origins to show that f1-ori ssDNA synthesis initiation and termination functions overlap, but can be inactivated separately by modifying distinct sequences (18). In 1992, Specthrie *et al*. packaged ssDNA as short as 292 bases into phage-like particles they called microphages (19). They were able to build small ssDNA strands using a phagemid that included an f1-ori, a packaging sequence (PS), and a truncated f1-ori that acts as a terminator (f1-oriΔ29). The terminator interrupts ssDNA synthesis of the full phagemid sequence, leading to packaging and export of only the region flanked by the ori and terminator.

**Figure 1.**
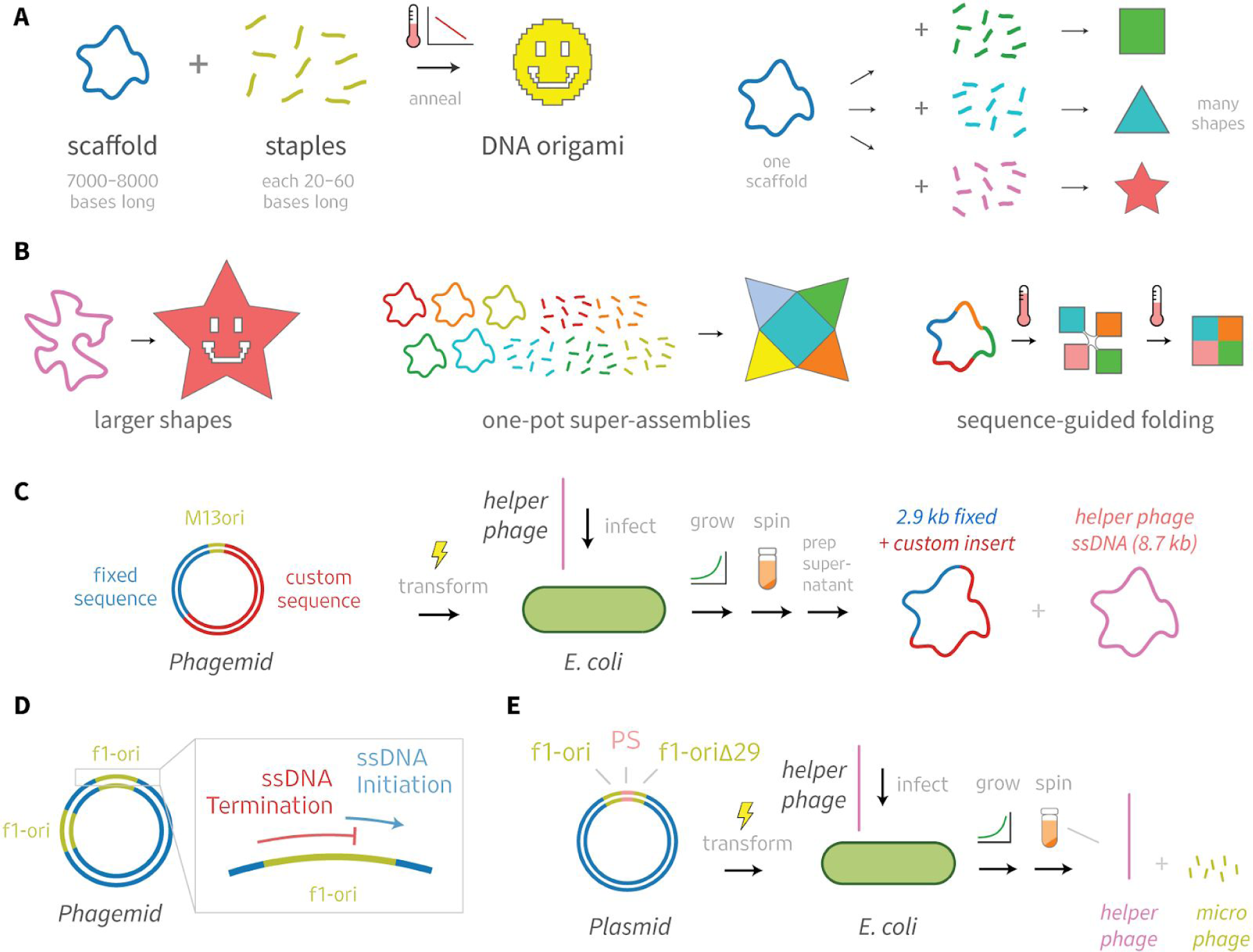
DNA origami design would benefit from custom scaffolds. **A** Many DNA origami shapes can be folded from a single scaffold. **B** New scaffolds will expand the space of possible designs. **C** Phagemids are excellent sources of scaffolds, but have multi-kilobase sequence constraints. **D, E** Previous studies offer hints of how these constraints could be circumvented. Dotto *et al*. (18) used phagemids with modified origins to show that f1-ori ssDNA initiation and termination functions overlap, but can be inactivated separately by modifying distinct sequences. Specthrie *et al*. (19) produced phage-like particles with ssDNA as short as 292 bases using a truncated f1-ori that acts as a terminator (f1-oriΔ29).

## Materials and Methods

### Construction of pScaf vector

We initially converted the pUC18 vector into a phagemid by cloning the M13 origin from M13mp18 at the *NdeI* and *KpnI* restriction sites. Digested fragments were ligated with T4 DNA ligase (NEB) and transformed into XL1-Blue MR competent cells (Agilent). We grew the pUC18-M13ori phagemid and recovered the DNA with a miniprep. Next, we used M13mp18 as a template to PCR-amplify the ssDNA synthesis terminator, based on the Δ29 design from Specthrie *et al. (19)* The terminator was then inserted into the *BamHI* and *EcoRI* sites of the pUC18-M13ori. All subsequent variants of the terminator (Fig 1) were assembled by PCR and cloned into the same *BamHI* and *EcoRI* sites, and verified for correctness using DNA sequencing. We used the variant with 3 thymine bases (TTT) as the final pScaf vector in all subsequent experiments in this work.

### Cloning scheme

We used *KpnI* and *BamHI* restriction sites between the ssDNA synthesis initiator and terminator regions to clone inserts into the pScaf vector (Figure 3). In one step we cloned short PCR-amplified insert sequences (A, B, C) directly into pScaf vectors that then generated the three smallest scaffolds (1512, 2268, and 3024). To create longer custom scaffolds, we adapted a multi-step cloning scheme previously used for cloning repeat protein cDNA constructs (25). We amplified custom sequence inserts in a polymerase chain reaction (PCR) using forward and reverse primers designed to incorporate into the product a 5’ *KpnI* and *BglII* site, and a 3’ *BamHI* site, respectively. We cloned PCR-amplified sequence inserts of lengths up to 2.5 kb into separate pScaf vectors (D–L). To combine two inserts to create one larger insert, we digested the pScaf vector containing the intended 5’ region with *PvuI* and *BamHI*, and digested the vector containing the intended 3’ region with *PvuI* and *BglII*. *BamHI* and *BglII* digestions create compatible sticky-end overhangs that can be used to ligate the two inserts. The bases adjacent to the compatible sticky ends are different, so the ligation product does not reconstitute an internal *BamHI* or *BglII* site. However, the ligated fragment maintains a 5’ *BglII* site and a 3’ *BamHI* site for subsequent cloning rounds. We digest each plasmid at the *PvuI* restriction site to split the ampicillin-resistance gene *bla* into two parts that are only reconstituted by successful ligation of the full-length product. Thus, we generated three vectors with tandem inserts (DE, FG, IJ), and then combined FG with H to create FGH. We combined IJ with K to create IJK, which was subsequently combined with L to create IJKL. The pScaf vectors containing inserts DE, FGH, and IJKL were then ready to produce the larger scaffolds (5544, 8064, and 10080).

### Scaffold amplification and purification

We transformed XL1-Blue-MRF’ using the M13cp helper plasmid (24) and then made it chemically competent with TSS (10% PEG-8000, 30 mM MgCl2, 5% DMSO, in 2xYT, pH 6, filtered) in order to produce a competent XL1-Blue_Helper strain. XL1-Blue_Helper was transformed with the desired pScaf construct to create a custom-phage-producing strain. We selected and grew a single colony for 18 hours in an incubator-shaker (30°C, 225 rpm) in 3-mL 2xYT media (1.6% tryptone, 1% yeast extract,.25% NaCl) containing kanamycin, carbenicllin, and chloramphenicol. Subsequently, the culture was transferred to a shake flask containing 100 mL 2xYT, 10 mL phosphate buffer (7% potassium phosphate dibasic, 3% sodium phosphate monobasic, pH 7, autoclaved), 1 mL 50% glucose, 0.5 mL 1 M MgCl_2_, and the appropriate antibiotics. The flask was then grown for 24 hours on a shaker (30°C, 225 rpm). The flask was harvested by transferring the culture to a 500-mL ultracentrifuge bottle, stored on ice for 30 minutes, and then centrifuged at 7000g for 15 minutes at 4°C, pelleting the bacteria while allowing the phage to remain in the supernatant. The phage-containing supernatant was transferred to a clean 500-mL ultracentrifuge bottle, where 4 g PEG-8000 and 3g NaCl were added. The bottle was then incubated on ice for 30 minutes prior to centrifugation at 9000g for 15 minutes at 4°C, allowing the phage to form a pellet. The supernatant waste was decanted and the phage pellet was resuspended in 3 mL TE buffer. The bottle was then centrifuged at 15000g for 15 minutes at 4°C to pellet any residual bacteria. The supernatant containing the concentrated phage was then transferred to a 50-mL ultracentrifuge tube. 6 mL lysis buffer (0.2M NaOH, 1% SDS) was added and the tube was mixed. Next, 4.5 mL neutralization buffer (3M KOAc, pH 5.5) was added and the tube was again mixed. The tube was incubated on ice for 15 minutes and then centrifuged at 15000g for 15 minutes at 4°C. The resulting supernatant was then decanted into a new 50-mL ultracentrifuge tube and 27 mL pure ethanol was added to it. The tube was capped and mixed by inversion prior to incubation at -20°C for 18 hours. After incubation, the tube was centrifuged at 16000g for 15 minutes at 4°C to pellet the DNA. The supernatant waste was decanted leaving the DNA pellet at the bottom of the tube. The pellet was washed with 9 mL ice-cold 70% ethanol and centrifuged at 16000g for 5 minutes at 4°C. The supernatant waste was again decanted. The pellet was subsequently dried by gently blowing air into the tube, and finally the dried pellet was resuspended in up to 1 mL TE, depending on the desired concentration.

### DNA origami preparation

Custom scaffold and staples were mixed at final concentrations of 20 nM and 200 nM, respectively, in a buffer containing 5 mM Tris, 1 mM EDTA and 13 mM MgCl2. The solution was then subject to the following temperature ramp: denaturation at 65°C for 15 minutes, followed by cooling from 60°C to 38°C with a decrease of 1°C per 50 minutes.

### Agarose gel analysis

Purified scaffolds and folded origami products were analyzed using 2% agarose gel electrophoresis in TBE (45 mM tris-borate and 1 mM EDTA) supplemented with 11 mM MgCl2 and SYBR Safe. Upon sample-loading, gels were run for 2 hours at 80V and subsequently scanned using a Typhoon FLA imager. DNA origami products folded from 1512-, 2268-, and 3024-base scaffolds were subsequently purified by extracting the desired gel band using a razor blade, muddled to break down the agarose, and then centrifuged through a Freeze ‘N Squeeze column.

### PEG purification

DNA origami products folded from scaffolds of 5544 bases and larger were purified using PEG precipitation. Assembly products were mixed with an equal volume of PEG precipitation buffer (15% PEG-8000, 10 mM Tris, 20 mM MgCl2, 500 mM NaCl) and immediately centrifuged at 16000g at 20°C for 25 minutes. The supernatant was removed and the pellet (which is often invisible) was resuspended in 1x folding buffer with magnesium (5 mM Tris, 1 mM EDTA and 13 mM MgCl2).

### Transmission electron microscopy

Purified origami structures were diluted to approximately 25 ng/μL prior to imaging. 5 μL of the diluted origami was applied to glow-discharged, carbon-coated, 400-mesh formvar grids (Ted Pella) for 1.5 minutes. The grid was then blotted dry on filter paper (Whatman). Washing and staining was performed by preparing a pierce of parafilm with two 15-μL droplets of 1x folding buffer and two 15-μL droplets of 2% aqueous uranyl formate stain solution. The grid was dipped onto the first buffer droplet, blotted dry, dipped onto the next buffer droplet, blotted dry, dipped onto the first stain droplet, blotted dry, and then held onto the final stain droplet for 45 seconds before being blotted dry. Grids were then allowed to air-dry for 10 minutes prior to imaging. Electron micrographs were collected using an FEI TECNAI T12 transmission electron microscope using a using a 4k × 4k charge-coupled device camera (UltraScan 4000, Gatan) at 26000× and 52000× magnifications. Class averages were obtained using EMAN2 software.

## Results

We constructed a phagemid, pScaf, that can be used to create custom DNA origami scaffolds (Figure 1). We converted pUC18 into a phagemid for custom ssDNA production by adding four components: a full-length M13 origin for ssDNA initiation, *KpnI* and *BamHI* restriction sites for insert cloning, the M13 packaging sequence for phage particle export, and a modified M13 origin to serve as the ssDNA synthesis terminator. When we tested the “Δ29” terminator used by Specthrie *et al.* (19), we were able to produce our desired custom ssDNA scaffolds. However, our phage cultures also yielded an off-target ssDNA species consistent in size with residual packaging of the region downstream of our custom insert. We suspected that the Δ29 terminator may not fully abolish initiation of ssDNA synthesis, resulting in multiple ssDNA species produced from a single phagemid similar to constructs reported by Dotto *et al.* (18). Thus, we sought to optimize the terminator to reduce spurious initiation and produce a more pure ssDNA scaffold product (Figure 2).

**Figure 2:**
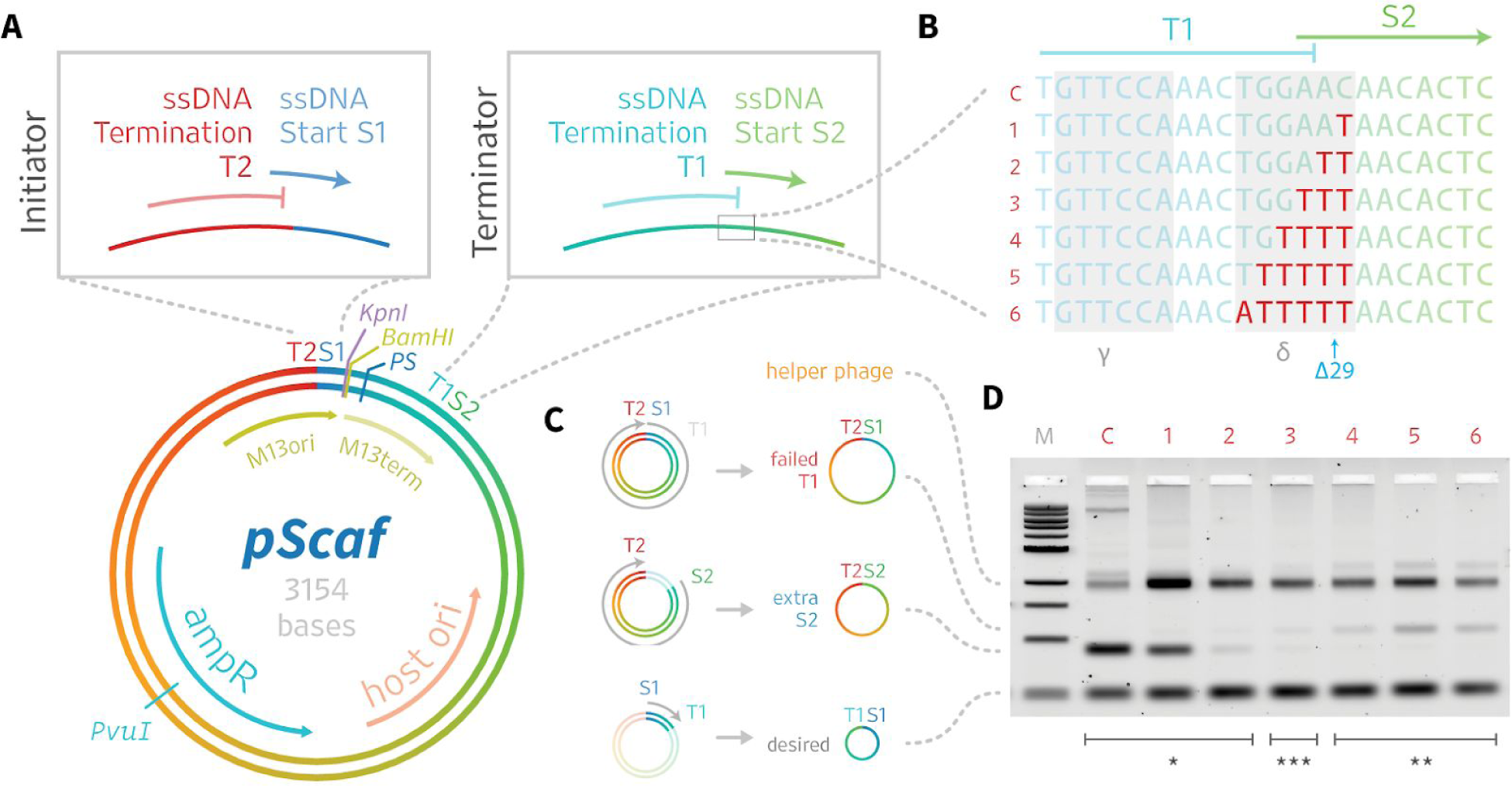
Construction and Optimization of the *pScaf* phagemid. **A** The pScaf phagemid is derived from the pUC18 plasmid and M13mp18 phage vectors. The phagemid includes an M13 origin of replication (M13ori) that contains ssDNA start site S1 and ssDNA termination site T2, a *KpnI* and *BamHI* cloning site, a packaging sequence (PS), and a terminator of ssDNA synthesis (M13term) containing ssDNA termination site T1 and ssDNA start site S2. The *M13term* sequence was adapted from the *M13ori* sequence by selectively deactivating the S2 ssDNA initiation function. **B** The ssDNA synthesis initiation and termination functions of the origin overlap. Therefore, we used a mutational screen of the δ region to optimize the M13term performance. **C** Three primary species were observed: S1T2, representing failed termination at terminator T1, S2T2, representing spurious initiation at initiator S2, and S1T1, the desired product. **D** We analyzed the variants by agarose gel. Substituting 0–2 thymines yielded the S2T2 species (*). Substituting 4–6 bases yielded the S1T2 species (**). Substituting 3 thymines yielded the best balance between spurious S2 initiation and failed T1 termination, producing the most pure S1T1 species (***).

### Terminator optimization

We started by truncating the last 50 bases of the 381-base M13 ori sequence so it included the core origin of plus-strand replication (*α,β, γ*, and *δ* regions) but omitted the downstream initiation enhancer region (26). We performed a mutational screen starting at the 3’ end of the δ region, near the site of the Specthrie *et al.* Δ29 truncation. We cloned variants of the phagemid with up to six δ-region bases substituted with thymines (T), except for one thymine base which we replaced with adenine (Figure 2c). We tested ssDNA production of each variant in *E.coli* culture co-infected with helper phage. We observed that 0-, 1-, or 2-base substitutions permitted ssDNA initiation, leading to synthesis of an off-target species of ssDNA. Meanwhile, 4-, 5-, or 6-base substitutions interfered with ssDNA termination, leading to an alternate off-target ssDNA species. We achieved the best balance between target and off-target ssDNA using the 3-base substitution, which resulted in relatively low amounts of each off-target species. We subsequently used the 3-base substitution in all pScaf constructs.

### Custom scaffolds and test origami

We created custom scaffolds with lengths of 1512, 2268, 3024, 5544, 8064, and 10080 bases (Figure 3). Each scaffold was verified by sequencing and analyzed by agarose-gel electrophoresis. We used Cadnano (27) to design a series of DNA origami shapes that approximate rectangular cuboids or brick-like structures. We folded and analyzed the resulting DNA origami shapes using agarose gel electrophoresis and negative-stain TEM (Figure 4).

**Figure 3:**
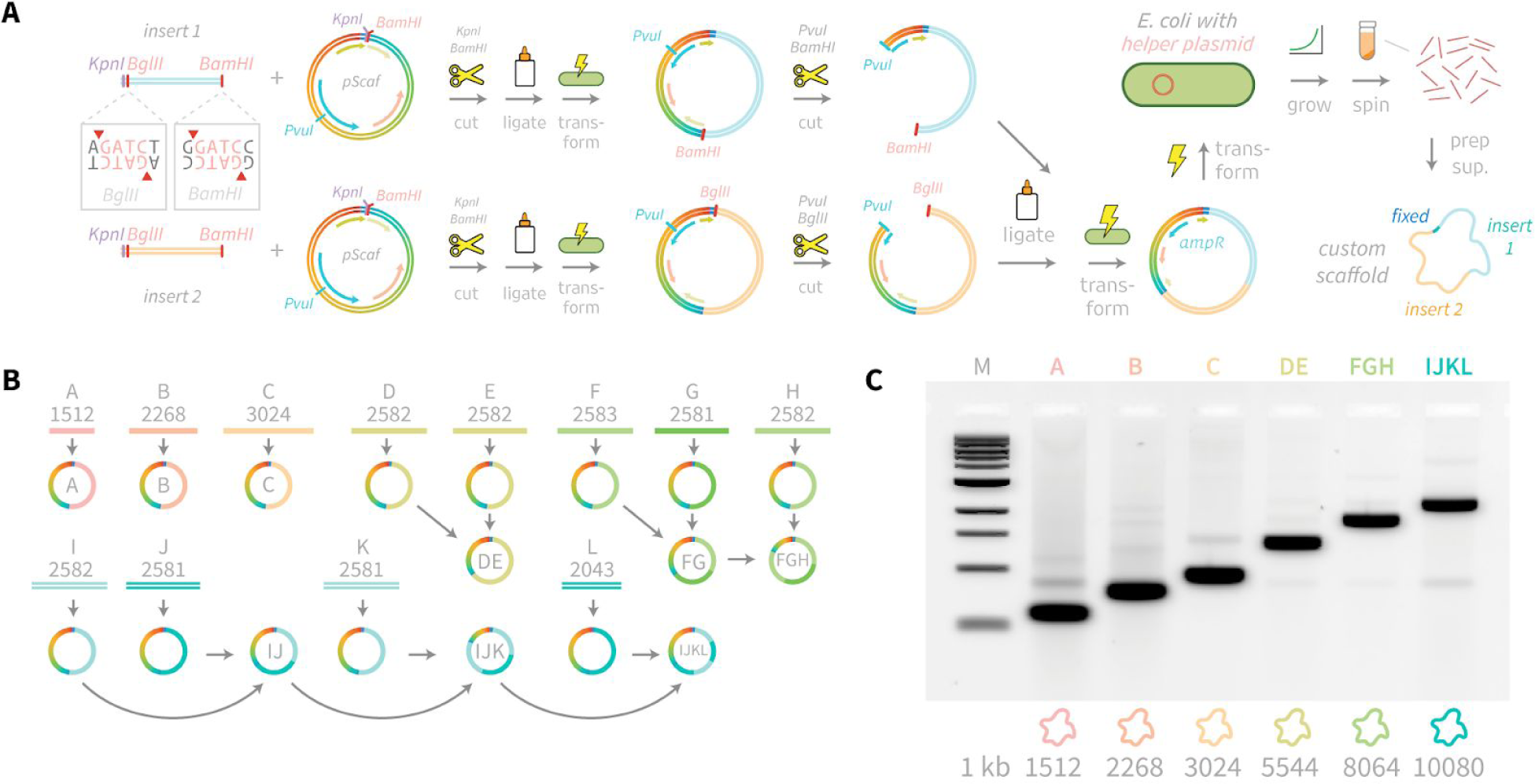
Cloning scheme and gel analysis of new scaffolds. **A** Custom sequence inserts are PCR-amplified with a forward primer containing *KpnI* and *BglII* sites and reverse primer containing a *BamHI* site. Inserts up to 3 kb in length were directly transformed into *E.coli* bearing helper plasmid. Larger scaffolds can be assembled by iterative *PvuI+BamHI* digestion of the vector containing the 5’ fragment, and *PvuI+BglII* digestion of the vector containing the 3’ fragment, followed by ligation, transformation, and miniprep. **B** Twelve inserts (A–L) were cloned into *pScaf* vector at the KpnI-BamHI site. Inserts A, B, and C were used to produce scaffolds with lengths of 1512, 2268, and 3024 bases. Larger scaffolds were assembled in multiple rounds as shown. **C** All scaffolds were grown in XL1-Blue cells containing helper plasmid M13cp, recovered and analyzed by agarose gel electrophoresis to determine purities ranging from 46% to 83%.

**Figure 4:**
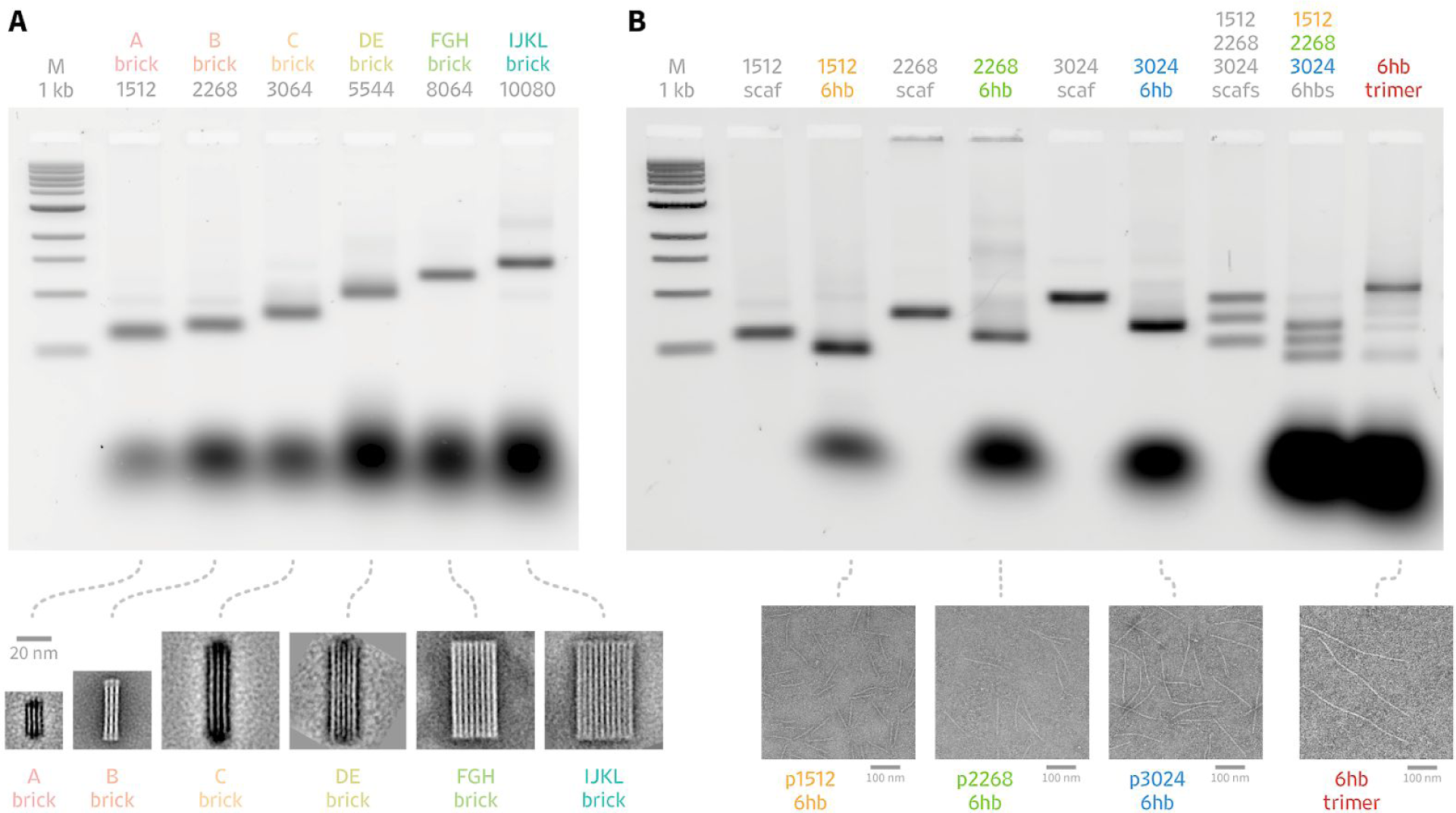
Folding Custom Scaffolds into DNA Origami Shapes and Assemblies. Six scaffolds of varying sizes were folded into DNA origami shapes and assemblies. Folding reactions were analyzed by gel electrophoresis, and origami analyzed negative stain transmission electron microscopy. **A** Brick-like DNA origami shapes were folded from scaffolds of lengths 1512, 2268, 3024, 5544, 8064, and 10080 bases. Each brick was PEG-purified and imaged by negative stain TEM. Class averages for each brick are shown, along with a 20-nm scale bar. **B** Scaffolds of lengths 1512, 2268, and 3024 were folded separately into six-helix-bundle (6hb) nanotubes. All three scaffolds and 6hb-staple sets were combined into one-pot reaction mixtures. When connector staples are included, the three scaffolds assemble into a 6hb trimer. TEM micrographs of the individual 6hb nanotubes and combined trimer are shown with 100-nm scale bars.

### One-pot folding of multi-scaffold assemblies

The ability to synthesize custom scaffolds with relatively little sequence overlap affords the possibility of folding DNA origami from multiple scaffolds in a one-pot reaction. To validate this concept, we designed a six-helix-bundle nanotube comprised of 3 scaffolds with lengths of 1512, 2268, and 3024 bases. The scaffolds were folded in a one-pot reaction both with and without connector staples to join them into a single nanotube.

## Discussion

Our approach has significant advantages over pre-existing methods for making custom scaffolds to build DNA origami nanostructures. First, we have demonstrated that custom scaffolds can be generated with lengths of up to 10 kb. Thus, the method is compatible with nearly all origami shapes that have been published to date, and will enable the creation of additional large shapes. Second, although cloning large (>3kb) custom insert sequences into phagemids presents difficult challenges, the cloning scheme we report here circumvents some of them. We found the insertion of long sequences is much more reliable with a multi-step assembly approach that cuts the antibiotic resistance gene during each round, and includes a positive selection for full-length constructs that reconstitute the gene. We typically only needed to screen a handful of colonies of transformed *E.coli* for each cloning round to locate the desired sequence. Third, because we rely on phagemids to grow custom scaffolds in *E.coli*, the method can be used to produce milligram-scale quantities of custom scaffolds using flasks and a shaker-incubator. Bacterial production offers a potentially reliable and feasible method for scaling up production to much larger quantities using bioreactors, as has been demonstrated for other scaffolds (20, 28).

With our method, it will now be possible to rapidly generate large DNA origami shapes with highly customized scaffold sequences. We recognized the need for easier methods to make custom scaffolds when making DNA origami shapes that fold from limited sets of reusable staple sequences (29). Here, we chose scaffold lengths to be multiples of 42, a convenient repeat length for designs using the honeycomb lattice. For convenience, we incorporated sequences from pre-existing plasmids such as pFastBac-GFP-dynein-2(D1091–Q4307). Additional scaffolds of various lengths can be generated quickly using a similar approach. Sequences also might be ordered from gene synthesis vendors and combined with our approach to generate scaffolds tailored for answering specific questions about DNA origami design principles, or to incorporate protein-binding sites.

Custom scaffolds will be useful for studying the DNA origami method itself. For example, new scaffolds can be used to examine sequence-level determinants of folding yield. Sequences of particular base compositions (high-GC, high-AT, 3-letter alphabets) with well-defined repetition or self-complementarity can be constructed and used to test hypotheses explaining the role of melting temperature or secondary structure in origami folding. Additionally, custom scaffolds can have practical benefits for routine origami design. We have often desired to create large DNA origami shapes with specific custom sequences at multiple non-contiguous locations within the scaffold. Generating these custom scaffolds is now much easier compared to cloning large inserts directly into M13mp18 in a single step.

It is also useful to be able to produce multiple unique scaffolds with very little sequence overlap. Sets of multiple unique custom sequences can be used for facile one-pot assembly of multimeric asymmetric structures. We demonstrated the one-pot assembly of small six-helix-bundle nanotube shapes, but expect our approach can extend to reliably making larger shapes. For example, it should now be possible to design three custom 10-kb scaffolds that can assemble into a 30kb origami in which nearly the entire multimeric structure is uniquely addressable and can incorporate modified staples.

Before we settled on the pScaf design, we attempted to generate custom scaffolds using a phagemid with a terminator based on the Δ29 truncation used by Specthrie *et al.* (19). Our observations of an off-target ssDNA species led us to attempt to optimize the terminator sequence. To guide our approach and select the site for our mutational screen, we reviewed previous studies of the initiation and termination of filamentous phage plus-strand synthesis (18, 26, 30). A better understanding of the fundamental biology of filamentous phages will likely lead to further improvements to the pScaf phagemid.

Although the TTT terminator variant greatly improved the scaffold purity, we were not able to completely eliminate incomplete termination or spurious initiation in our scaffold preps. We also observed some prep-to-prep variation in scaffold quality and production of off-target species when using the helper plasmid. We analyzed the scaffold gel image with ImageJ and estimated that the purity of correct-length scaffold products ranged from 46% to 83% of total integrated intensity of the sample lane (Fig 3c). Our maximum purity is similar to commercially available M13-derived scaffold (31). Our DNA origami gel analysis and TEM images show that we obtained a relatively homogeneous population of folded origami structures, so the current levels of purity will be suitable for many applications. Future optimization of the ori and terminator sequences, helper plasmid, host strain, or growth conditions may yield cleaner target ssDNA products.

The sequence insertion locus used *KpnI* and *BamHI* restriction sites, which would need modification if those sites are to be incorporated into a custom scaffold. A multiple cloning site would provide greater flexibility for cloning sequence inserts. We did not attempt to reduce the size of the terminator region that is incorporated into the scaffold ssDNA, so a further reduction in size of the fixed region of 381 bases may be possible with additional optimization. While we produced scaffolds up to 10 kb in length, we note that this length does not represent an upper limit. We have not yet attempted to make longer scaffolds, but it should be possible following our scheme.

In summary, we have developed and validated a novel phagemid, pScaf, for creating highly customized ssDNA DNA origami scaffolds of 10-kb lengths that can be produced at milligram-scale yields. Our approach removes a long-standing constraint that has held back progress in the use of large DNA origami scaffolds, which is that researchers have had to rely on the genome of M13 phage which cannot be easily customized. The ability to generate many long, unique custom scaffolds will enable researchers to resolve many unanswered questions about what the optimal methods are to create and design DNA origami sequences, how to easily incorporate functional sequences, such as protein-binding sites, into multiple sites within each shape, and how to better adapt these methods in future applications.

## Acknowledgements

We thank P. Rothemund and L. Bienen for comments on the manuscript. We are grateful to A. Bradbury for providing the M13cp helper plasmid. This work was supported by the Army Research Office (W911NF-14-1-0507), Del E. Webb Foundation (13-2-28), and National Science Foundation (CCF-1317640). T.A. holds a Ruth L. Kirschstein NRSA Postdoctoral Fellowship (F32GM119322). S.M.D holds a Career Award at the Scientific Interface from the Burroughs Wellcome Fund (1010247).

## Author Contributions

P.M.N., T.A., and S.M.D. design the research. P.M.N. performed the experiments. P.M.N. and T.A. collected and analyzed the TEM data. All authors contributed to data interpretation and to writing the manuscript.

